# Multiple ecological and evolutionary mechanisms drive treatment-induced antibiotic resistance

**DOI:** 10.64898/2026.02.18.705331

**Authors:** Matthew J Shepherd, Niamh E Harrington, Anastasia Kotarra, Claudia Igler, Kenny Cagney, Taoran Fu, Elizabeth M Grimsey, Joanne L Fothergill, Dylan Z Childs, Steve Paterson, Michael A Brockhurst

**Affiliations:** Division of Evolution, Infection and Genomic Sciences, School of Biological Sciences, University of Manchester, M13 9NT, UK; Institute of Infection, Veterinary and Ecological Sciences, University of Liverpool, L69 3BX, UK; Department of Animal and Plant Sciences, University of Sheffield, S10 2TN Sheffield, UK

## Abstract

The emergence of resistance within patients during antibiotic treatment is an important cause of treatment failure. However, the ecological and evolutionary mechanisms driving within-patient emergence remain poorly understood. Here, we analysed 24,478 *Pseudomonas aeruginosa* isolates sampled from 180 bronchiectasis patients during a clinical trial to understand how ciprofloxacin-resistant infections emerged over a one-year period of pulse-dosing. Pre-existing resistance predominated, accelerating resistance emergence relative to patients where resistance emerged by spontaneous mutation or strain immigration. Selective sweeps of costly mutations increased resistance over time in some patients, whereas in others oscillating resistance levels were driven by antibiotic treatment and resistance-growth trade-offs between genetically divergent subpopulations. Our findings show that infections under identical treatment follow diverse and sometimes complex ecological and evolutionary paths to antibiotic resistance, with implications for better predicting and managing treatment-induced antibiotic resistance.

## Introduction

Rising rates of antibiotic resistance threaten global health, endanger modern medical treatments, and cause millions of deaths annually (*1, 2*). The molecular mechanisms of antibiotic resistance and their incidence among bacterial genomes have been well described (*3, 4*), as has the transmission of resistant strains between humans, animals and natural environments (*5, 6*). Far less understood, however, are the ecological and evolutionary mechanisms by which antibiotic resistance emerges within human infections during antibiotic treatment. A recent review of observational clinical studies outlined the four main mechanisms causing antibiotic resistant infections (*7*): selection for pre-existing resistant genotypes, spontaneous gain of new resistance mutations, horizontal gene transfer of resistance genes from the surrounding microbiota, or immigration of resistant strains. Such clinical studies often involve low numbers of patients and limited pathogen sampling, so the relative importance of these mechanisms is unknown for most human infections. This limits our ability to avoid resistance through evolution-informed treatments (*8*), that could both improve patient outcomes and antimicrobial stewardship. An important exception are *Escherichia coli* urinary tract infections, where immigration of resistant strains from the patient’s microbiota predominates (*9*), enabling prediction of more effective treatments, but whether this pattern translates to other pathogens and infection types is unknown. Evolutionary theory and *in vitro* experiments suggest that the spread of antibiotic resistance can be limited by fitness costs in the absence of antibiotics (*10*). However, fitness costs of antibiotic resistance are rarely measured in clinical studies and their importance for driving *in vivo* resistance dynamics remains unclear. To understand the ecological and evolutionary mechanisms driving the emergence of antibiotic resistance, we performed a large-scale retrospective evolutionary analysis of *Pseudomonas aeruginosa* populations collected during a phase-III clinical trial for ciprofloxacin. *P. aeruginosa* is a common cause of chronic and difficult-to-treat infections (*11*) often associated with treatment-induced resistance (*12*), as such these infections are ideal for understanding the ecological and evolutionary mechanisms of within-patient resistance emergence.

## Results and Discussion

We analysed *P. aeruginosa* isolates from populations originally sampled from 180 bronchiectasis patients with established lung infections (*13*), who were randomly assigned to either treatment (n = 119) or placebo (n = 61) groups for the ORBIT-3 trial (*14*). Treatment group patients received cyclical ciprofloxacin dosing, comprising 28 days treated with inhaled liposomal ciprofloxacin (on-phase) followed by 28 days without ciprofloxacin (off-phase) for 6 cycles (also referred to as pulse-dosing), whereas placebo patients received empty liposomes. At the end of the trial, all patients from both the treatment and placebo groups were given ciprofloxacin for 28 days during an Open Label Extension (OLE) phase. For our study, we sampled *P. aeruginosa* infections most intensively (n = 90 random colonies per patient) at baseline prior to treatment to enable robust detection of rare pre-existing resistance. For subsequent timepoints, during both on-phases and off-phases of treatment cycles, we sampled 15 colonies per patient, shown to be sufficient to representatively capture population diversity from sputum samples (*15*) (see Fig. S1 for study design). This yielded 24,478 *P. aeruginosa* isolates (n = 16,177 from baseline; n = 8,301 from subsequent timepoints). For every isolate we quantified ciprofloxacin minimum inhibitory concentration on Iso-Sensitest agar (MIC; Fig. S3) and maximal growth rate in King’s B nutrient broth (Fig. S4). Overall, ciprofloxacin MIC remained stable over time in the placebo group, whereas in the treatment group ciprofloxacin MIC increased during on-phases (Estimate = 2.01 µg/mL, SE = 0.55, df = 460, t = 3.65, p < 0.001, linear mixed-effects model (LMM)) then decreased during off-phases (Estimate = - 1.760 µg/mL, SE = 0.345, df = 460, t = -5.096, p < 0.001; contrast test on estimated marginal means (EMMs), Fig. 1A), confirming that treatment selected for ciprofloxacin resistance. Among the 86 treatment group patients for whom samples from both baseline and subsequent timepoints were available, breakpoint resistance, defined here by the breakpoint ≥4 μg/mL used in the ORBIT-3 trial (*14*), emerged in 52 (60.5%), with the remaining 34 (39.5%) not attaining breakpoint resistance at any point during the trial (full per-patient breakdown of each evolutionary mechanism assignment and evidence provided in supplementary spreadsheet file E1). However, the temporal dynamics of resistance during ciprofloxacin treatment varied markedly between patients. Infections grouped into 3 distinct trajectories of ciprofloxacin MIC dynamics: For 33 treatment group patients where breakpoint resistance emerged and time-courses were adequately sampled (Online Methods), we observed stable (n=8), monotonic (n=7), or oscillatory (n=18) resistance dynamics (Fig. 1B).

**Figure 1:**
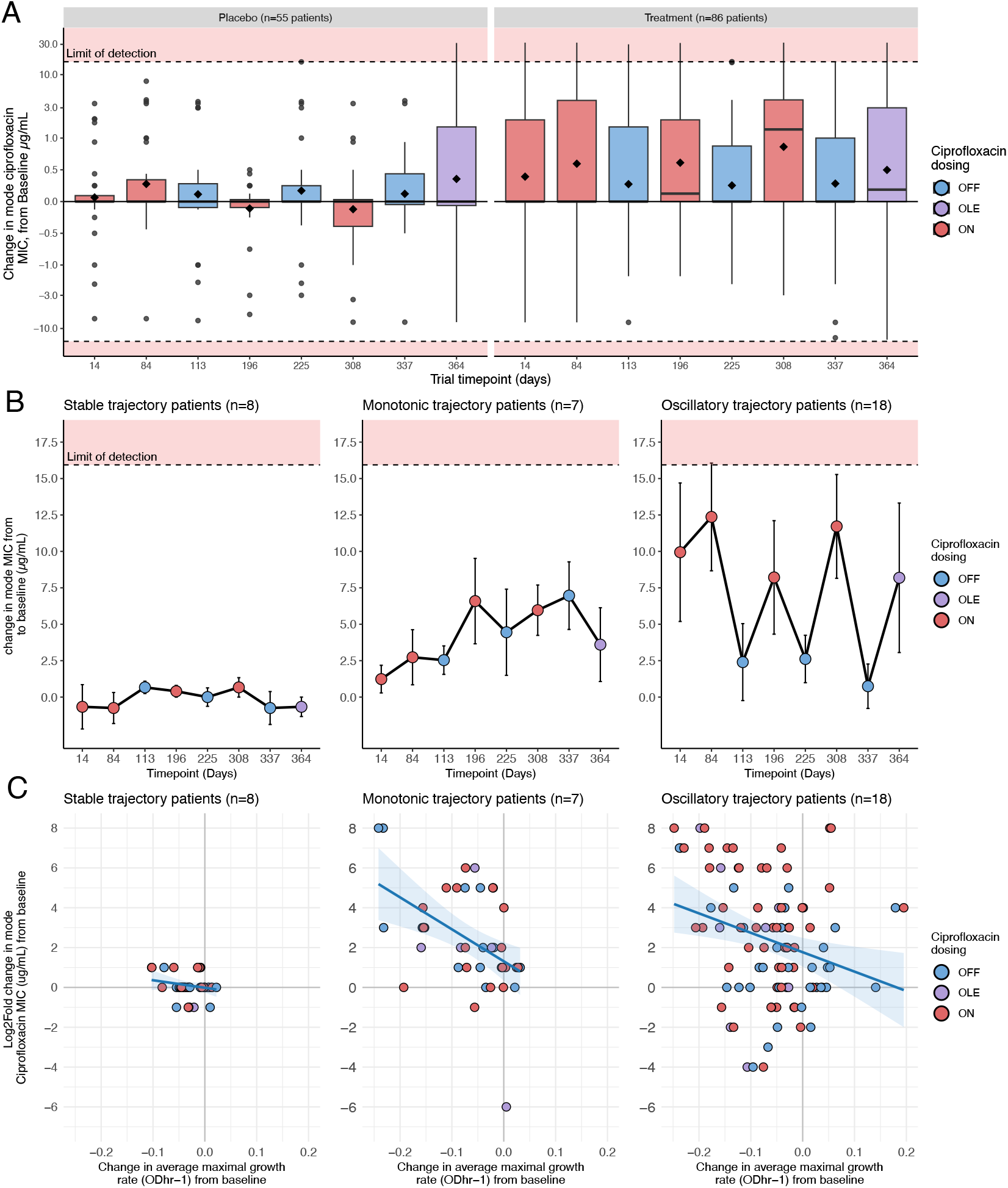
Ciprofloxacin minimum inhibitory concentration (MIC) dynamics over time and its relationship with growth rate. In all sub-plots, treatment phase indicated by colour: on-phase = red; off-phase = blue; open label extension = purple. The limit of detection of the MIC assay is indicated by dashed lines with red shaded areas. **A)** Change in mode ciprofloxacin MIC relative to baseline over time. Boxes indicate interquartile range for the change in mode ciprofloxacin MIC relative to baseline mode MIC (μg/mL) and whiskers show range, points denote outliers. **B)** Change in mode ciprofloxacin MIC relative to baseline over time per resistance trajectory class. Points show change in mode MIC relative to baseline ± standard error. **C)** Relationship between resistance and growth rate per resistance trajectory class. Points show mean values of all isolates per patient per sampling point. Trendlines give linear regression fits and the ribbons show 95% confidence interval.

Consistent with fitness costs of gaining ciprofloxacin resistance, increased resistance was significantly associated with reduced growth rate for monotonic (Linear model (LM), slope = - 0.0068, SE = 0.0023, t =-2.92, p = 0.0057) and oscillatory (slope = -0.0033, SE = 0.0012, t = - 2.78, p = 0.0067) trajectories, but not for stable trajectories (slope = -0.00035, SE = 0.0019 t = - 0.18, p = 0.86). For oscillatory trajectories only, we also observed significant clustering by treatment phase (PERMANOVA, 999 permutations, Euclidean distance, F(1,95) = 10.10, R^2^ = 0.096, p = 0.002). As such, oscillatory trajectories alternated between two states, higher resistance with slower versus lower resistance with faster growth, driven by antibiotic selection on a growth-resistance fitness trade-off.

We next identified the genetic mechanisms underlying observed changes in ciprofloxacin resistance. We sequenced genomic DNA pools of all isolates per sample for all samples, as well as whole-genome-sequenced 4206 individual clones from baseline (2857; n = 16 per sample, (*13*)) and subsequent timepoints (1349; n = 4 per sample with highest ciprofloxacin MIC). We identified variants in genes known to be associated with ciprofloxacin resistance (*16*), principally: *gyrA* or *gyrB* encoding DNA gyrase; *parC* or *parE* encoding topoisomerase IV; *mexR* or *nfxB* encoding regulators of multidrug efflux, and *mexAB-oprM, mexCD-oprJ*, or *mexEF-oprN* efflux pumps. We also performed multi-locus sequence typing and quantified core genome SNP variation to enable detection of immigrating strains or the presence of coexisting divergent subpopulations. Combining these genomic datasets with MIC data allowed us to pinpoint genetic variants associated with the initial emergence of breakpoint resistance (Online Methods explain the details of how ecological and evolutionary mechanisms were assigned; Fig S2). Breakpoint resistance arose through three, non-exclusive primary mechanisms within the treatment group, showcasing the complexity of resistance evolution even under single antibiotic treatment (Fig. 2A). Selection for pre-existing resistance was the predominant mechanism (50% of patients with resistance, n = 26), followed by spontaneous mutation (25% of patients with resistance, n = 13), and strain immigration as the least common mechanism (11.5% of patients with resistance, n = 6). For a further group of patients, gain of resistance could not be linked to a causal genetic variant (unclear mechanism, 19.2% of patients, n = 10). We never observed gain of resistance through horizontal gene transfer, potentially due to the predominance of point mutations as mechanistic causes of ciprofloxacin resistance with only limited mobile resistance genes (*17*). These results contrast those in acute *E. coli* urinary tract infections where strain immigration predominates, including for ciprofloxacin (*9*), indicating that the relative importance of mechanisms of resistance emergence likely varies with body site and pathogen species.

**Figure 2:**
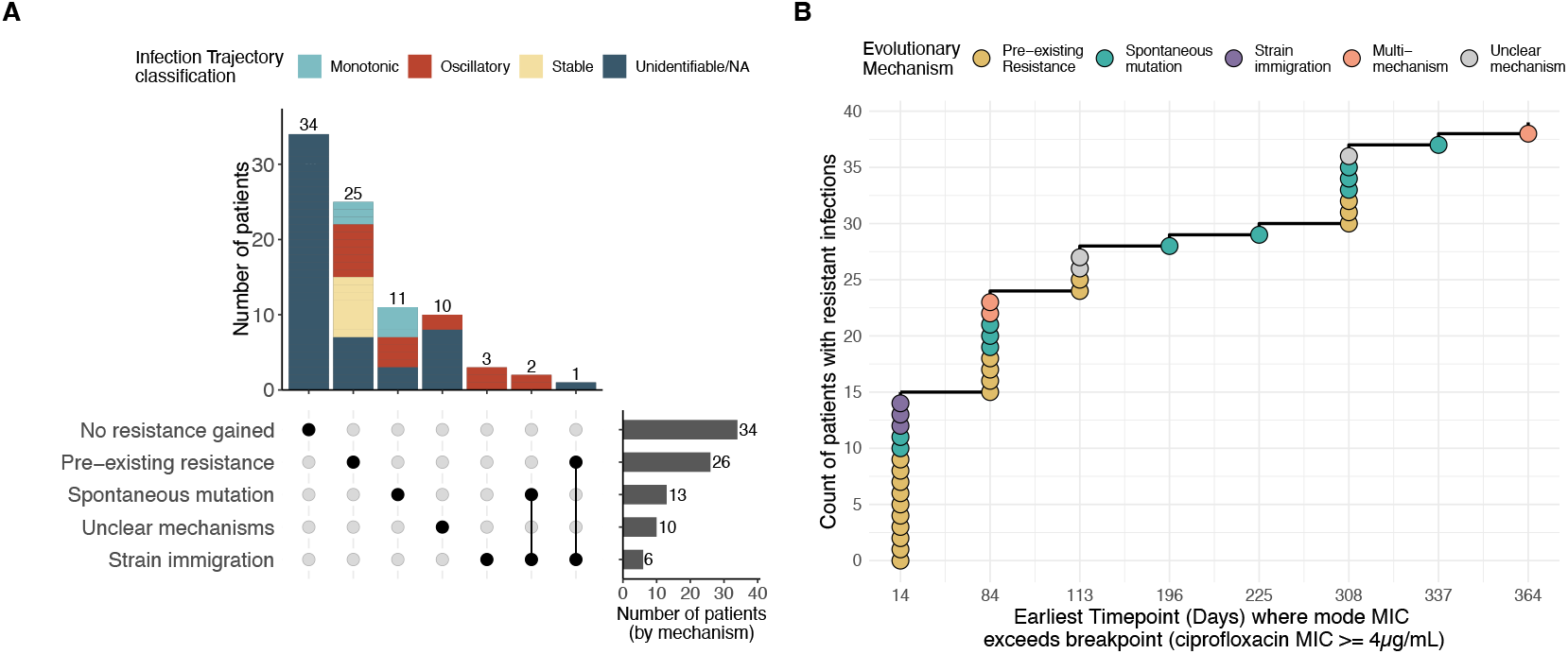
Ecological and evolutionary mechanisms of resistance emergence within-patients. **A)** Upset plot of within patient resistance emergence mechanisms (n= 86 patients). Bold points in dot matrix denote mechanisms and linked points indicate combinations of multiple mechanisms. Right hand bar plot indicates total number of patients per emergence mechanism. Upper stacked bar plot indicates number of patients for each emergence mechanism grouping, with colours indicating proportions of patients per MIC trajectory class (known for n = 33 patients; see legend). **B)** The speed of resistance emergence varies by mechanism. Time-to-event plot counting the number of patients with mode MIC greater than breakpoint per sampling point. Points are coloured by emergence mechanism (see legend).

The dominance of pre-existing resistance in our study is concerning, because it accelerated resistance emergence compared to all other mechanisms (χ^2^(1) = 18.5, p < 0.001; Fig. 2B). Most of the infections where pre-existing resistance drove initial emergence had attained breakpoint resistance by the first subsequent sampling point on day 14 (i.e., mid-point of the first on-phase, Fig. 2B). In principle, emergence by pre-existing resistance could be easier to combat clinically, through more intensive pre-treatment sampling of the infecting population to detect resistant genotypes, enabling more accurate diagnosis and improved prescription and stewardship of antibiotics (*18*). In contrast, emergence of breakpoint resistance driven by spontaneous mutation or strain replacement was less predictable, being distributed over the duration of the trial, as expected for these inherently stochastic population processes. Consistent with evolutionary theory, therefore, these data show that adaptation from standing genetic variation is faster and more deterministic than mutation or gene flow-driven adaptive processes (*19*).

There was not a direct correspondence between MIC trajectories and resistance emergence mechanisms. Although stable trajectories were exclusively associated with pre-existing resistance, both monotonic and oscillatory trajectories could arise through multiple emergence mechanisms (Fig. 2A, S5). Importantly, however, whilst initial emergence of breakpoint resistance was attributable to single mechanisms in most patients, over the course of the trial more complex combinations of mechanisms contributed to MIC dynamics in many infections (Fig. S6). Typically, these cases involved gain of additional spontaneous resistance mutations by already resistant lineages, further increasing their MIC. As such, it is likely detrimental to continue to treat resistant infections with the same antibiotic, because evolution of resistance continues to progress after breakpoint resistance is gained.

We next asked if different resistance trajectories corresponded to different underlying allele frequency dynamics. Allele frequency dynamics at resistance loci were predictive of MIC trajectory in 77% of patients (Fig. 3A; CI = (76%, 96%), p = 0.034, non-parametric bootstrap, 1000 replicates, two-sided test, H_0_ = 1/3), and varied with phase between MIC trajectory classes (Fig. 3B; LMM, type III ANOVA, phase × trajectory interaction; χ^2^(2) = 121.96, p < 0.001). Stable trajectories showed almost no change in resistance loci allele frequencies, consistent with resistance being pre-existing prior to treatment in all these patients. Monotonic trajectories could also arise from pre-existing resistance but were more commonly associated with spontaneous mutations (Fisher’s Exact Test, OR = 6.76, 95% CI: 0.81–67.27, p = 0.042), which frequently underwent selective sweeps (Fig. S7). Monotonic trajectories were thus characterised by strong positive selection of resistance loci alleles during on-phases and weak or absent negative selection during off-phases (Fig. 3B, Fig. S8, Contrast Test on EMMs from LMM on-phase versus off-phase: estimate = 0.076, SE = 0.018, z = 4.03, p < 0.001; Wald test versus zero: on-phase, estimate = 0.071, SE = 0.033, z = 2.16, p = 0.03; off-phase, estimate = -0.005, SE = 0.034, z = -0.15, p = 0.88). Regardless of trajectory, spontaneous resistance mutations were often costly (Fig. S9A median change in average maximal growth rate < 0, p = 0.001, Wilcoxon test), and for multiple monotonic trajectories costly resistance stably persisted following a selective sweep despite periods of relaxed antibiotic pressure under pulse dosing (Fig. S9B, S9C). As such, persistence of resistance *in vivo* is not a reliable proxy for an absence of fitness cost as previously suggested based upon genetic data alone (*20*). For oscillatory trajectories, resistance emerged by pre-existing resistance (n= 7), spontaneous mutation (n= 4), strain immigration (n= 3), combinations of these (n= 2), or unclear mechanisms (n= 2) across different patients. Regardless of this diversity of emergence mechanisms, resistance oscillations were characterised by strong positive selection on resistance loci alleles during on-phases and strong negative selection during off-phases (Fig. 3B, Contrast Test on EMMs from LMM on-phase versus off-phase: estimate = 0.22, SE = 0.0096, z = 22.87, p < 0.001; Wald test versus zero: on-phase, estimate = 0.081, SE = 0.019, z = 4.23, p <0.001; off-phase, estimate = -0.14, SE = 0.02, z = -6.88, p <0.001), consistent with the resistance-growth fitness trade-off we observed in these patients (Fig. 1C). Leveraging such fitness trade-offs to promote reinvasion of susceptible genotypes between treatment phases may explain the clinical benefit of pulse-dosing (*21*) and could improve treatment if predictable (*22*).

**Figure 3:**
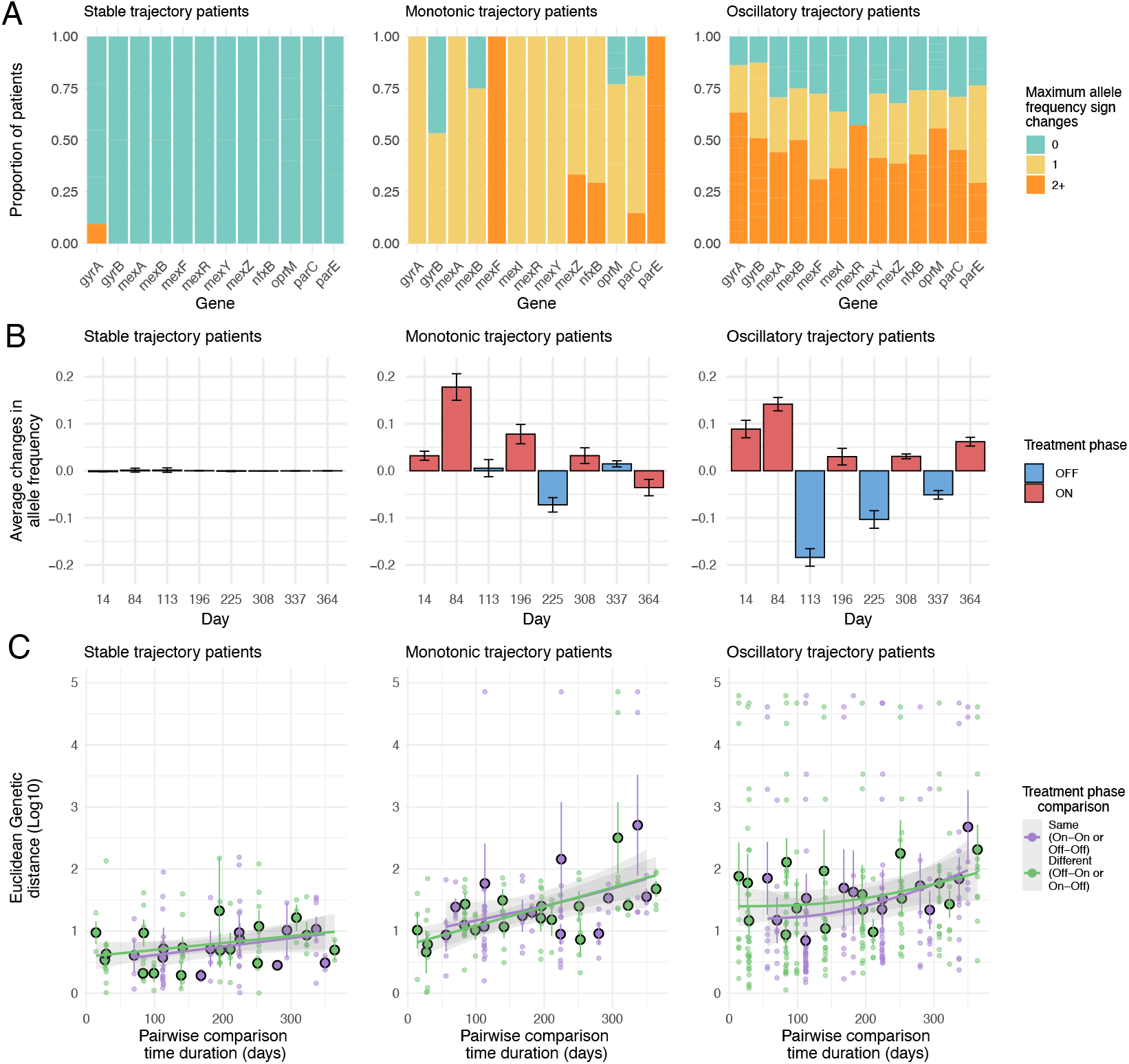
Resistance trajectory classes are associated with distinct allele frequency dynamics at resistance loci and contrasting patterns of genome-wide pairwise genetic distance. **A)** Proportions of patients per maximum number of allele frequency sign changes per resistance locus. Colours denote maximum number of allele frequency sign changes (Green = no sign change; yellow = one sign change; orange = two or more sign changes. **B)** Changes in resistance loci allele frequency per sampling period per MIC trajectory class. Bars show mean change in allele frequency at resistance loci ± standard error. Treatment phase is indicated by colour (red = on phase; blue = off phase; purple = open label extension phase (OLE)). **C)** Euclidean genetic distance between all pairs of samples per patient plotted against time difference. Large dots show mean genetic distance ± standard error; small dots show genetic distance for each pairwise comparison. The trendlines show fits from generalised additive model and the ribbon denotes 95% confidence interval. The treatment phase comparison (i.e., whether the pair of samples are from the same or different treatment phases) is indicated by colour (same phase = purple; different phase = green).

We hypothesised that rapid oscillations of resistance would require ecological alternation of distinct resistant and susceptible subpopulations with treatment phase, rather than recurrent evolutionary transitions. To test this, we analysed genome-wide pairwise genetic distances between all samples within a patient. Consistent with ecological alternation between subpopulations, we observed higher genetic distances between pairs of samples from different (i.e., on/off) versus the same (i.e., on/on or off/off) treatment phases for oscillatory (Fig. 3C; Contrast Test on EMMs from LMM, parametric bootstrap, r = 1000: estimate = 35.1, 95% CI 6.29 – 62.01, p = 0.022) but not stable (estimate = 6.67, 95% CI -39.95 – 52.01, p = 0.77) or monotonic (estimate = -2.96, 95% CI -43.18– 38.93, p = 0.89) trajectories. Together, these data suggest that different resistance trajectories were associated with distinct underlying allele frequency dynamics. Notably, whereas monotonic trajectories typically featured selective sweeps of resistance alleles, oscillatory trajectories more often featured fluctuating resistance allele dynamics (Fig. S7). Which trajectory arose appeared linked to whether the infection contained genetically divergent subpopulations: Where present, relatively resistant versus susceptible subpopulations could alternate, driven by selection on a resistance-growth trade-off. Where absent, populations progressively gained costly resistance. Thus, competition between ecologically coexisting subpopulations may be critical for leveraging fitness trade-offs to select against resistance within patients (*23*).

## Conclusion

Here, we provide an unprecedented understanding of the ecological and evolutionary dynamics of antibiotic-induced resistance during a clinical trial. In patients infected by the same pathogen and receiving identical antibiotic treatment, we observed multiple, varied and sometimes complex ecological and evolutionary mechanisms driving resistance emergence. Mechanisms varied in their speed and predictability, with pre-existing resistance notably accelerating resistance emergence compared to spontaneous mutation or strain immigration, whilst also being highly predictable and thus more easily diagnosable pre-treatment. We show fitness costs of resistance were common and drove oscillating resistance in phase with antibiotic selection where distinct resistant and susceptible subpopulations coexisted within a patient, highlighting where such trade-offs could be leveraged to reduce resistance. Otherwise, fitness costs posed little barrier to resistance evolving and persisting long-term. Indeed, continuing antibiotic treatment after resistance emerged in such patients could drive the further evolution of high-level resistance by sweeps of additional spontaneous mutations within already resistant lineages, suggesting that switching antibiotic would be of benefit both for the patient and antibiotic stewardship. Together, our findings highlight multiple ways that antibiotic diagnosis and treatment could be improved by integrating knowledge of the ecological and evolutionary processes occurring within treated patients to guide more personalised and robust antimicrobial therapies.

## Acknowledgements

The University of Dundee provided access to samples used in this research under a collaboration agreement. We are grateful to the patients, all hospital and clinical trial staff, and all others involved in ORBIT-3 trial. Genome sequencing was performed by the Centre for Genomics Research (CGR) at the University of Liverpool. We acknowledge Joel Doherty for assistance with image processing and James Chalmers for providing access to lab facilities. This work was supported by NIHR Manchester Biomedical Research Centre (NIHR203308). The project was funded by a grant from the Wellcome Trust (220243/Z/20/Z). CI is supported by a Wellcome Trust Early Career Award (225565/Z/22/Z).

## Authorship CRediT (Contribution Roles Taxonomy) statement

**Matthew J Shepherd:** Conceptualization, Methodology, Software, Formal analysis, investigation, Data curation, Writing - Original Draft, Writing - Review and Editing, Visualisation

**Niamh E Harrington:** Conceptualization, Methodology, Software, Formal analysis, Investigation, Data curation, Writing - review & editing, Visualisation

**Anastasia Kotarra:** Conceptualization, Methodology, Investigation, Data curation, Writing - Review & Editing

**Kenny Cagney:** Methodology, Investigation, Writing - Review & Editing

**Taoran Fu:** Methodology, Software, Writing - Review & Editing

**Claudia Igler:** Methodology, Formal analysis, Writing - Review & Editing, Visualisation, Funding acquisition

**Elizabeth M Grimsey:** Methodology, Investigation, Writing – Review and Editing

**Joanne L Fothergill:** Conceptualization, Writing - Review & Editing, Supervision

**Dylan Z Childs:** Conceptualization, Methodology, Software, Formal analysis, Writing – review and editing, Supervision, Funding acquisition

**Steve Paterson:** Conceptualization, Methodology, Software, Formal analysis, Visualization, Writing - Review & Editing, Project administration, Supervision, Funding acquisition

**Michael A Brockhurst:** Conceptualization, Methodology, Visualization, Writing – original draft, Writing - Review & Editing, Project administration, Supervision, Funding acquisition

