## Supplemental text and figures for "Multiple ecological and evolutionary mechanisms drive treatment-induced antibiotic resistance"

### Supplement

### Methods

#### Trial study design, clinical samples and bacterial isolation

The ORBIT-3 clinical trial was a one-year randomised, placebo-controlled phase-III clinical trial of an inhaled liposomal ciprofloxacin therapy for treatment of patients with the chronic lung condition bronchiectasis, and bacterial lung infections of the pathogen *Pseudomonas aeruginosa* (*1*). The trial involved a pre-treatment screening phase where patients were included based on presence of at least one ciprofloxacin sensitive *P. aeruginosa* isolate (a ciprofloxacin minimum inhibitory concentration <4 μg/mL). Patients were then dosed with aerosolised liposomal ciprofloxacin (or empty liposomes for the placebo group) via a nebuliser in a phased 28 ‘on’ and 28 ‘off’ cycling treatment regimen, for a total of six cycles (Fig. S1). After these six cycles totalling 336 days, a final 28 day ‘open label extension’ (OLE) phase was included, during which all patients including the placebo group were given active treatment with ciprofloxacin. Previously (*2*), sputum samples were taken from 180 patients during the screening phase (referred to as ‘baseline’), and a total of 16,177 *P. aeruginosa* isolates collected (~90 per patient, providing a high degree of internal biological replication for each sample).

In this work, we isolated *P. aeruginosa* from sputum samples taken at up to eight timepoints during the dosing period of the trial (referred to as ‘timecourse'). These timepoints were at days 14, 84, 113, 196, 225, 308, 337 and 364 of the trial (Fig. S1). In total, we were able to isolate 8301 *P. aeruginosa* colonies (15 per timepoint, per patient, again providing a high degree of internal biological replication for each sample), with 141 of the 180 patients sampled at baseline having at least one timepoint sample during the timecourse. Bacterial isolation was achieved as described previously (*2*); in brief, sputum was treated with Sputasol (SR0233 Oxoid), and colonies isolated on selective Cetrimide Agar (22470, NutriSelect® Plus). Isolates were further cultured in King’s B medium (KB, recipe detailed below in phenotypic tests subsection) in 96-well plate format, and stored at -80°C. Isolation of *P. aeruginosa* was confirmed by species-specific 16S rRNA gene PCR (detailed in (*2*)).

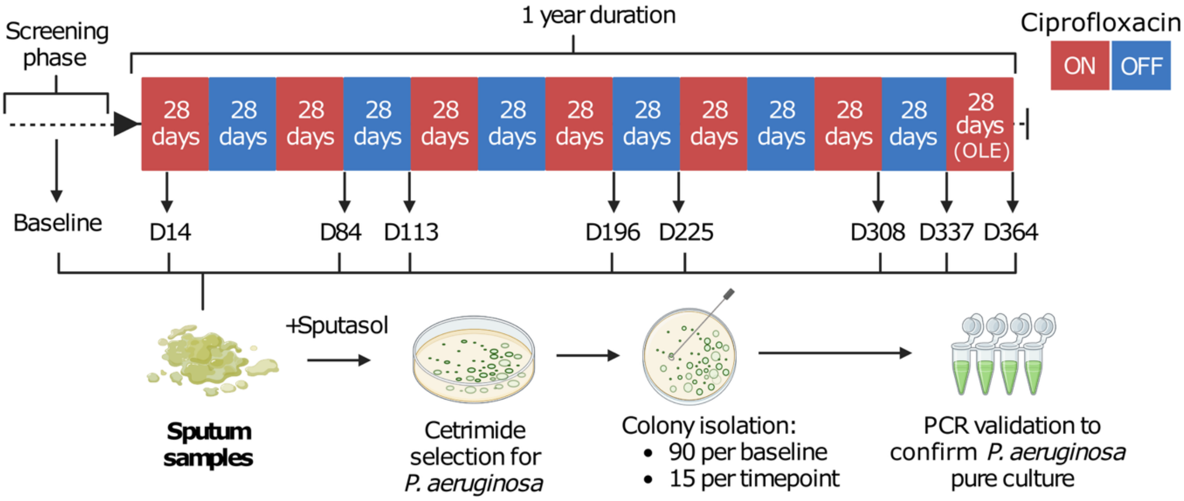


**Figure S1:** Schematic of ORBIT-3 trial dosing regimen. Coloured bars indicate dosing, with red denoting ‘on’ phases where patients received the study drug/placebo, and blue denoting 'off' phases where treatment was paused. Timepoints where sputum was sampled for *P. aeruginosa* isolates are indicated below the coloured bars.

**Bacterial growth rate assay**

For all 24,478 *P. aeruginosa* isolates we tested susceptibility to ciprofloxacin, and bacterial growth curve dynamics. To achieve this in a high-throughput manner, isolates were first cultured in KB broth (20 g $l^{-1}$ Bacto proteose peptone No.3 (Gibco), 1.5 g $l^{-1}$ Potassium phosphate dibasic trihydrate (P5504, Sigma), 1.5 g $l^{-1}$ Magnesium sulfate heptahydrate (M1880, Sigma) and 10 g $l^{-1}$ Glycerol (49770, Honeywell)) and 96-well format, alongside control *P. aeruginosa* reference strains PAO1, LESB58, and ATCC27853, for ~40 hours static at 37°C. Cultures were used to inoculate fresh KB broth at a final inoculum dilution of 1 in 150 in a 96-well plate. Plates were sealed with Breath-Easy® membrane seals (Diversified Biotech) to allow aeration of the cultures during incubation. Plates were incubated at 37°C in an Agilent BioTek LogPhase 600 Microbiology Reader with orbital shaking (800 rpm), and optical density at 600 nm (OD600) was recorded at 20-minute intervals. For every isolate, two technical replicate growth curves were measured. Bacterial maximal growth rates were calculated for every isolate. Raw OD600 values were log-transformed using the natural logarithm (ln), and the maximal growth rate was calculated as the largest slope of the log-transformed OD600 across three consecutive timepoints. This approach reduced spurious results due to sudden OD600 fluctuations between individual timepoints, which are common in growth curve data. The maximal growth rates from both technical replicates were then averaged to obtain a single representative value for each isolate.

**Ciprofloxacin susceptibility testing**

To test ciprofloxacin susceptibility, plates of Iso-Sensitest agar (Oxoid) were prepared supplemented with ciprofloxacin. Ciprofloxacin stocks were prepared in solution of 1% acetic acid and Milli-Q ultrapure water, and used to produce agar plates at the following final concentrations of ciprofloxacin (all in μg/mL): 0, 0.0625, 0.125, 0.25, 0.5, 1, 2, 4, 8, 16. Bacterial cultures were serially stamped onto these plates in the order listed, using a URI®*DOT* multipoint inoculator (MAST) with 2.4mm inoculum pins. Plates were incubated for 16 hours at 37°C, and images recorded of all plates using a PhenoBooth+ precision imager (Singer Instruments).

To determine the ciprofloxacin minimum inhibitory concentration (MIC), we developed an image analysis script in MATLAB (available at https://github.com/TaoranFu/PIA). Colony sizes were first extracted from images after background and light reflection correction. Growth was confirmed on the ciprofloxacin-free control plate (a colony size larger than 4 $mm^2$), and the reduction in colony size relative to this control was determined for plates containing ciprofloxacin. A reduction in colony size greater than 95% was taken as indication of significant growth inhibition. The MIC was then recorded as the smallest concentration of drug which achieved this reduction. Significant growth in the highest concentration (16 μg/mL) was recorded as an MIC over limit of detection (>16), and failure to grow on the lowest concentration of ciprofloxacin tested was recorded as under limit of detection (<0.0625).

**DNA extraction, whole genome sequencing and isolate pool sequencing**

We selected 4 *P. aeruginosa* isolates with the highest ciprofloxacin MIC for each timepoint and patient for whole genome sequencing. If more than 4 isolates shared the highest MIC, 4 were randomly chosen from that group. If fewer than 4 were available, isolates with the next highest MIC were randomly sampled. DNA was extracted and libraries prepared for Illumina sequencing as described previously (*2*), in brief; DNA was extracted using the Quick-DNA Fecal/Soil Microbe kit (Zymo Research), and concentration standardised using a Qubit Flex Fluorometer (Thermo Fisher Scientific). Library preparation was performed using NEBNext Ultra II FS DNA Library Prep Kit for Illumina (New England Biolabs), and libraries were amplified and 10bp index sequences (Integrated DNA Technologies) incorporated by PCR. Libraries were quantified and size analysis performed, followed by pooling in an equimolar manner, and concentrated using AMPure XP beads (Beckman Coulter). Average length and concentration of final libraries were measured, and samples were then sequenced on an Illumina NovaSeq 6000 150bp paired-end run by the Centre for Genomics Research, University of Liverpool.

Additionally, we performed sequencing on pools of *P. aeruginosa* isolates. Cultures of all isolates at each timepoint and patient were pooled (90 isolates per patient baseline, 15 isolates per timepoint for each patient), DNA extracted and sequenced as described above.

**Bioinformatic analysis**

Whole genome sequences for baseline isolates (2854 total) were collected and analysed previously (*2*), including multi-locus sequence type (MLST) analysis and core SNP calling (Sequences available through the European Nucleotide Archive (ENA): accession PRJEB65845). We used the same bioinformatics pipeline and packages for analysis of genomes collected for this work, in brief; reads were quality checked, *de novo* assembly performed (median contig number = 78) and genomes annotated. These were then used to perform MLST analysis to assign sequence types to isolates, and core SNP calling. Four reference strain genomes were used for these analyses: *P. aeruginosa* PAO1 (NCBI, GCF_000006765.1), *P. aeruginosa* PA14 (NCBI, GCF_000014625.1), *P. aeruginosa* LESB58 (NCBI, GCF_000026645.1) and *P. aeruginosa* PA7 (NCBI, GCF_000017205.1). All gene information was sourced from pseudomonas.com (*3*).

Reads from pooled sequencing were mapped using bowtie2 (v 2.4.2) against either PAO1 or PA14 (NCBI, GCF_000006765.1, GCF_000014625.1, respectively) appropriate to the phylogroup type of each baseline sample and to a set of reference genomes from potential contaminants (*Acinetobacter baumannii*, *Achromobacter insuavis, Achromobacter piechaudii, Achromobacter ruhlandii, Achromobacter xylosoxidans, Delftia acidovorans, Delftia tsuruhatensis, Escherichia coli, Klebsiella pneumoniae, Morganella morganii, Stenotrophomonas maltophilia, Serratia marcescens*). Bam files containing reads mapping to the PAO1/PA14 reference were merged using samtools (v1.22.1) for samples within a patient (retaining sample information in the read groups) and freebayes (v1.3.6) used to count reference and alternate alleles with the –pooled-continuous option. Annotation for each SNP was added using snpEff (v5.2). R (v4.3.3) was then used to filter for a minumum depth of at least 10 reads in any sample within a patient supporting the reference or alternate allele and to cross-check and filter potential SNPs in the pooled sequencing against those found within single isolate sequences from that patient. Allele frequencies were then calculated for each sample within a patient and then SNPs retained that exhibited a frequency of >5% in at least one sample. Allele frequencies of specific resistance mutations were also extracted for separate analyses and visualisation.

**Isolate and sample *Pseudomonas aeruginosa* quality control steps**

To ensure all data analysed was from *Pseudomonas aeruginosa* only a set of quality control steps were implemented. As described above, isolates were screened with a set of *P. aeruginosa* specific 16S primers. Isolates which failed this screen were excluded. Despite this step, a number of isolates were identified as non-*Pseudomonas* only after whole genome sequencing (55 baseline isolates, 59 timecourse isolates). These isolates were excluded. Additionally, if all whole genome sequenced isolates from a specific timepoint were identified as non-*Pseudomonas,* the entire timepoint was also excluded (excludes timepoints that are likely to all be non-*Pseudomonas).* This removed 11 timepoints across 11 unique patients.

**Determination of evolutionary mechanism of resistance emergence**

Utilising the genotypic and phenotypic datasets we collected, we identified evolutionary mechanisms associated with resistant MICs (≥4 μg/mL) in the treatment group of the trial. For each patient, whole genome sequences of 4 pure isolates with highest MICs at each timepoint sample were compared to the 16 pre-treatment baseline isolate sequences. We focused on mutations / resistance alleles for the following genes known to be associated with ciprofloxacin resistance in *P. aeruginosa*; *gyrA, gyrB, parC, parE, nfxB, mexA, mexB, mexC, mexD, mexR, mexZ, oprM, and oprJ*. We sought to identify the evolutionary mechanisms driving emergence of any isolates with above-breakpoint MICs, even if this was only a minority of isolates, and/or transient, so evolutionary mechanisms were assigned to each patient on the basis of explaining how resistant isolates emerged within-patient. This was possible for 86 out of 119 treatment group patients, where we had a baseline sample and at least one timecourse sample.

Evolutionary mechanisms were assigned for each patient as follows (Fig. S2): Firstly, all patients where the MIC never crossed breakpoint at any timepoint sampled were assigned as ‘No resistance gained’. Then, for patients with a resistant MIC in their baseline sample, we checked if this MIC and the associated resistant genotype was maintained in at least one timecourse sample. If this was the case the patient was assigned ‘Pre-existing resistance’. Next, if resistant MICs were present in the timecourse only, we checked if the resistant isolates were of a different sequence type (ST) to the baseline sample, and that any new STs were not present in the baseline pool sequences for that patient. If so, and resistance alleles could be identified in the new ST, and no other conditions were met, the patient was assigned ‘Strain replacement’. Then, if resistant MICs were present in the timecourse and the ST remained the same as baseline, we checked for gain of mutations in resistance-associated genes. If these were found, we confirmed that these alleles were not present in the baseline pool sequences, and, if so, the patient was assigned ‘Spontaneous mutation’. If multiple of the above mechanisms were confirmed in a single patient, that patient was assigned as ‘Multi-mechanism’. Finally, if there were resistant isolates, for which an origin could not be ascribed to any of the above mechanism classifications, the patient was assigned as ‘Unclear’.

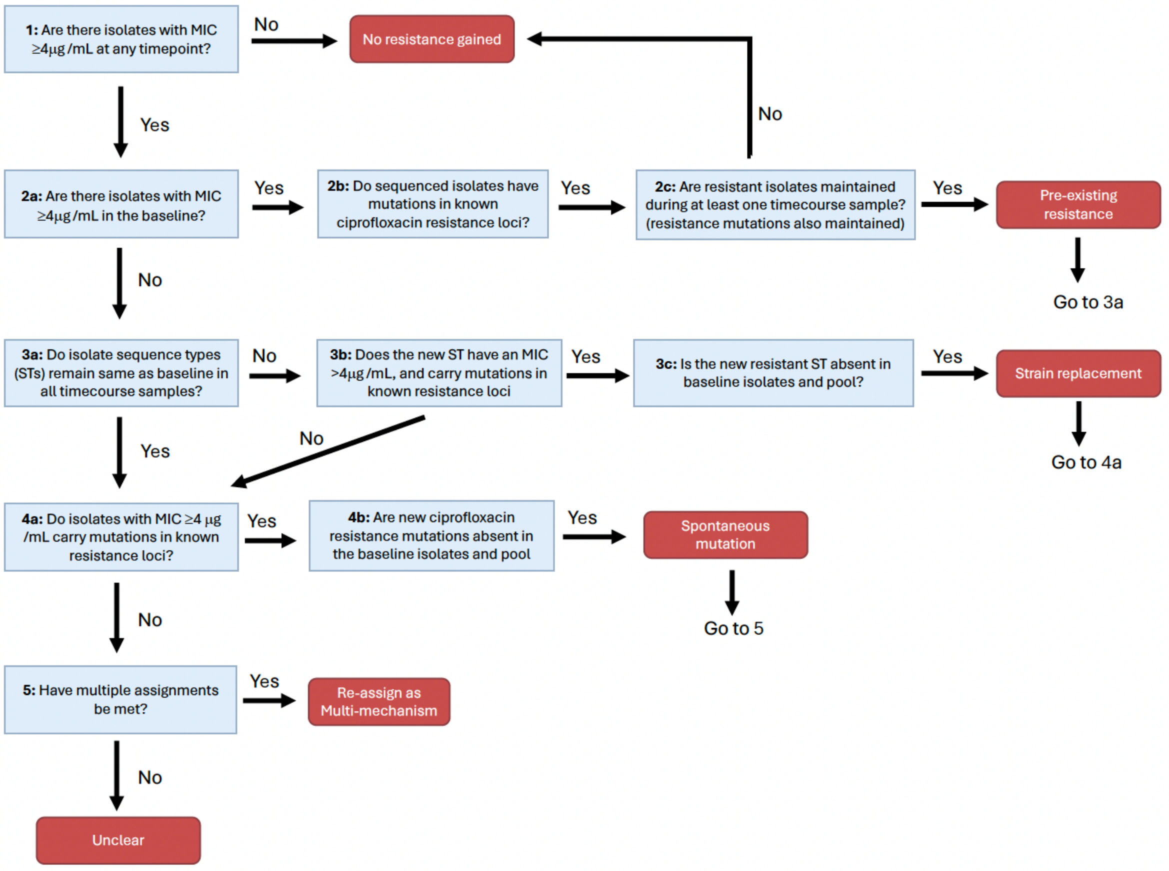


**Figure S2**: Flowchart for patient assignments of evolutionary mechanism of ciprofloxacin resistance emergence during the ORBIT-3 trial.

### Assignment of phenotypic trajectories of resistance emergence

Alongside evolutionary mechanisms, patients within which resistance emerged (i.e. a mode MIC ≥4 µg/mL was present) were assigned phenotypic trajectories based on how the mode MIC of the infection changed over time in response to treatment cycles. To do so, patients were required to have at least 3 timepoints sampled: the baseline pre-treatment sample, and 2 timecourse samples, including at least one on-phase and one off-phase. This was possible a total of 33 of the 52 patients where resistance emerged. Changes in mode MIC between each timepoint were smoothed so that only substantive shifts (>=2 Log2Fold changes in mode MIC) were considered, and the sign of changes determined (positive or negative). If no substantive changes in mode MIC occurred, the infection was assigned as displaying a 'stable' trajectory. Next, if there are 1 or more substantive changes in MIC that also involved sign changes (positive to negative or vice versa), the infection was assigned as 'cyclical'. Finally, if there were substantive changes in MIC that involved no sign changes, the infection was assigned as 'monotonic'. All 33 patients could be categorised into one of these three groupings.

### Statistical analyses and data handling

All data was handled and statistical analyses performed using R (version 4.2.2). For statistical analysis of MIC data, readings that sat outside assay limits of detection (>16µg/mL and <0.0625µg/mL) were set to the upper and lower bounds (16µg/mL and 0.0625µg/mL, respectively). This adjusted 359 values (1.47%) from >16μg/mL to 16μg/mL, and 460 values (1.88%) from <0.0625μg/mL to 0.0625μg/mL. This adjustment affected a small proportion of values (3.35% of the total), preserving rank ordering while compressing variation at the assay extremes, and is therefore expected to have minimal impact on the statistical analyses used in this study (detailed below), which rely on rank-based, permutation-based, or mixed-effects methods that are robust to non-normality and tied values. To understand overall MIC changes across the trial, we modelled the change in the mode MIC using a linear mixed-effects model (LMM, made using lmer from lme4 package) suitable for handling our large datasets which feature nested repeated measures. The model had the following structure: 'Change in mode MIC from baseline' ~ 'Trial group' * 'Treatment phase' + ( 1 | Patient). To determine whether ciprofloxacin MIC declined during off-phases for the treatment group after rising during on-phases, pairwise comparison of estimated marginal means from the above LMM (emmeans package), with a Benjamini-Hochberg correction.

Wilcoxon tests to establish differences between groups and Fisher's exact tests were performed using base R, which are suitable for handling non-normal, ordinal and low-n datasets. To determine the change in maximal growth rate – change in MIC correlation across trajectory classes, linear models were fit for each trajectory class separately, using base R. allowed direct comparison of association between changes in resistance and growth within each trajectory class whilst accounting for differences in intercepts across trajectories. PERMANOVA test for establishing trial phase clustering performed using the 'adonis2' function (999 permutations, vegan package), allowing for multivariate comparisons without assuming multivariate normality.

To investigate the rate of resistance emergence within-patient, we calculated the earliest timepoint where the mode ciprofloxacin MIC exceeds the breakpoint for each patient (note: this was possible for 39 out of 52 patients where resistance was detected (75%), as in the remaining 25% resistant isolates emerged, but remained in a minority of isolates so the mode MIC remained below breakpoint). A Kaplan-Meier curve was constructed for this data (survival package) grouping evolutionary mechanism into 'pre-existing' and 'other' (all other mechanisms). Differences in time to emergence of resistance between these two groupings was tested using a log-rank test (survdiff function from survival).

**Analysis of allele frequency dynamics**

*Processing of resistance allele frequency dataset*

To investigate resistance allele frequency dynamics, we filtered a full allele frequency dataset to only include non-synonymous variants in the following ciprofloxacin resistance-associated loci: *gyrA, gyrB, parC, parE, mexA, mexB, mexF, mexI, mexR, mexY, mexZ, nfxB, oprM, oprJ*. The dataset was further filtered to only include variants that appear in a ciprofloxacin resistant isolate genome (MIC ≥4 μg/mL), but also to exclude variants that are also present in a ciprofloxacin sensitive isolate from the same patient.

*Dynamics across the trial*

We determined patterns in allele frequency dynamics in the 31 patients with assigned phenotypic resistance trajectories based on MIC. To do so, allele frequencies were first discretized to binary values (0 or 1). This simplification was justified by the observation that most alleles showed rapid sweeps between fixation and loss over the trial period (Fig. 3B). For each patient, we counted the total number of discrete allele frequency transitions (from 0 to 1 or 1 to 0) between consecutive sampling time points at all resistance loci.

We then assigned each patient a trajectory category based on the maximal observed number of allele frequency transitions at any resistance locus. Trajectories were defined as 'stable' (0 changes), 'monotonic' (1 change), or 'cyclical' (2 or more changes). We calculated the predictiveness of allele frequency changes as the proportion of patients for whom trajectory assignments based on MIC and allele frequencies dynamics agreed. We determined confidence intervals and p-values for our ability to predict trajectories by using 1000 bootstrap replicates in R (functions *boot boot.ci* and *boot.pval*) and applying a two-sided hypothesis test with the null hypothesis that assignment to trajectories was random (i.e., 1/3).

We visualized the association between allele frequency changes and trajectory types, by aggregating the maximal observed number of allele frequency transitions across patients for each resistance locus.

*Dynamics across treatment phases*

We quantified absolute allele frequency changes for each trajectory across treatment phases by using only data points from consecutive sampling days (except for Day 84, which was calculated from Day 1 on to keep it consistent with other on-off-on phases), resulting in 3 off phases, 2 on phases and 3 mixed (on-off-on) phases (Fig. S2). As mixed phases contained more on- than off-phase time, and sampling occurred following an on phase, we assumed that on-phase effects dominated at these points (on-mixed).

For each trajectory type, we calculated the mean and standard deviation of absolute (sign-ignorant) allele frequency changes across all patients and resistance loci for the on, off and on-mixed phases. To assess differences in allele frequency dynamics between trajectory types and treatment phases, we applied a linear mixed-effects model (LMM) with phase, time, trajectory and their interactions as fixed effects, and patient and resistance locus as random effects. To determine how allele frequency changes varied with combinations of trial phase and trajectory, we used pairwise comparison of estimated marginal means from the above LMM (emmeans package) with a Benjamini-Hochberg correction, and to test if allele frequency changes differed from zero for each trial phase/trajectory combination, we used a Wald test on estimated marginal means (emmeans package). These tests allow direct comparisons between levels whilst accounting for the random effects structure of the LMM.

*Genetic distance analysis*

To investigate the driving forces behind different adaptive trajectory classes, we calculated the Euclidean genetic distance based on allele frequency data for all genes, and all pairwise comparisons of trial timepoints for each patient. The type of trial phase comparison (same = on vs on or off vs off ciprofloxacin treatment, different = on vs off), and the time span between each pair of compared timepoints was calculated. We then analysed this data using an LMM, and determined p-values using 1000 bootstrap replicates (non-parametric bootstrap), providing robust uncertainty estimates without assuming normality.

**Materials availability statement:**

All data and analyses supporting the results of this study are made publicly available where possible. All whole genome sequencing data (reads and assemblies) are available at European Nucleotide Archive (ENA) accession PRJEB65845. All phenotypic data and analysis code are available on GitHub (code used in MIC image analysis available here https://github.com/TaoranFu/PIA, code for all phenotypic analyses, trajectory assignments, and main test statistical tests available here https://github.com/mattshepherd502/within-patient-AMR-evo). Datasets for phenotypic and allele frequency analyses will be made available on the associated Zenodo record, DOI: 10.5281/zenodo.18482540

### Supplementary results

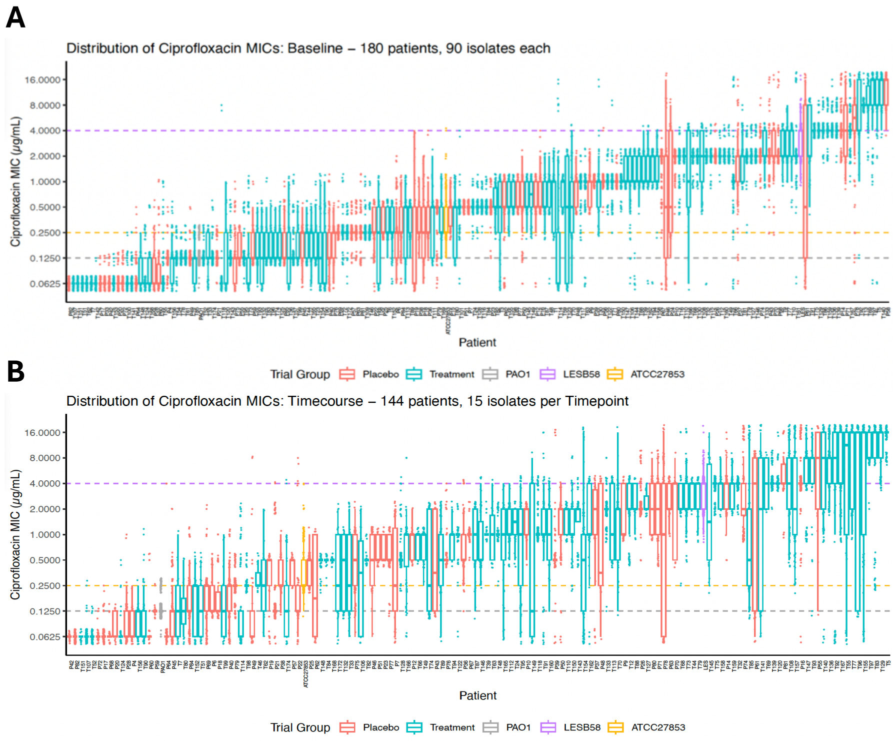


**Figure S3:** Distribution of isolate ciprofloxacin MICs (µg/mL), for A) baseline isolates, B) timecourse isolates. Each boxplot and data jitter represents a patient, with patient IDs listed along the x-axis. Boxplot boxes indicate the interquartile range. A data jitter is positioned beneath each boxplot. Colours indicate trial grouping of each patient (Red = placebo, blue = treatment), and data for three *P. aeruginosa* reference strains (Grey = PAO1, Purple = LESB58, Yellow = ATCC27853).

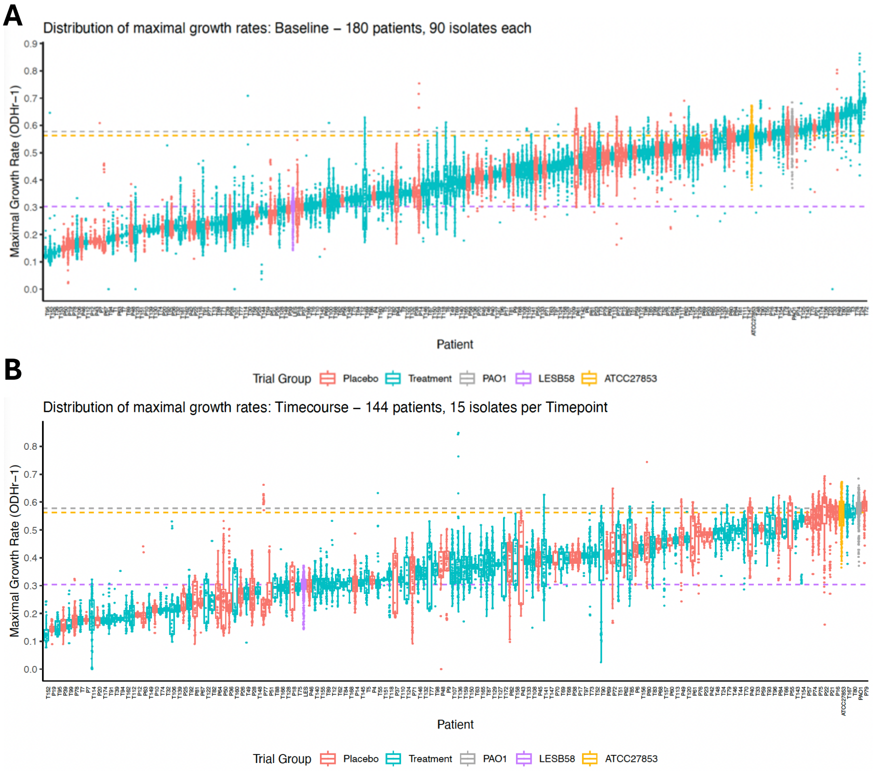


**Figure S4:** Distribution of isolate maximal growth rates (OD/hr), for A) baseline isolates, B) timecourse isolates. Each boxplot and data jitter represents a patient, with patient IDs listed along the x-axis. Boxplot boxes indicate the interquartile range. A data jitter is positioned beneath each boxplot. Colours indicate trial grouping of each patient (Red = placebo, blue = treatment), and data for three *P. aeruginosa* reference strains (Grey = PAO1, Purple = LESB58, Yellow = ATCC27853).

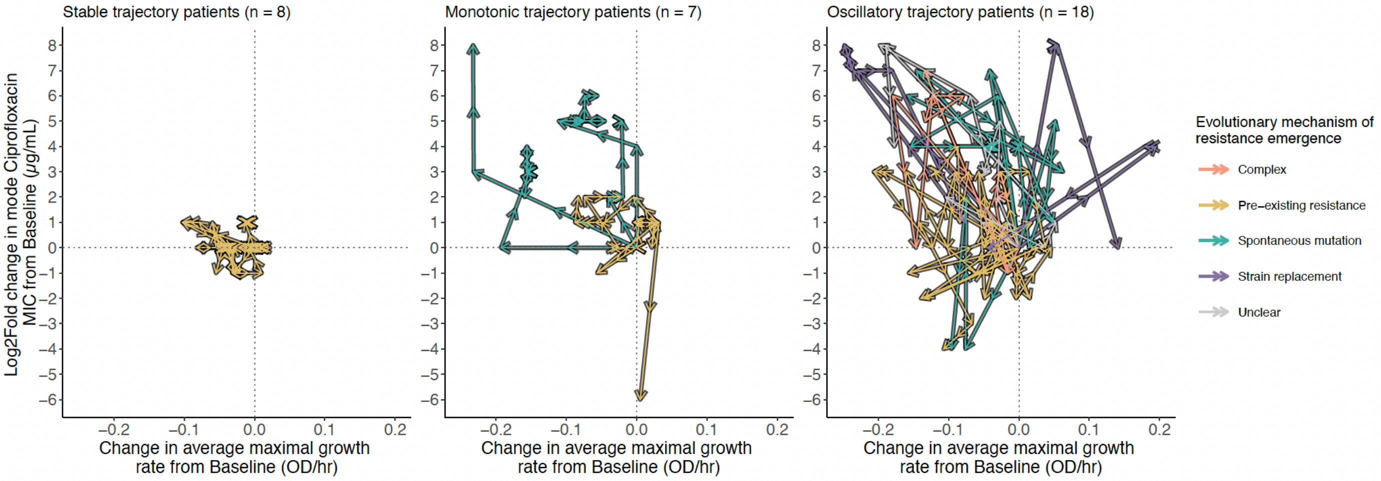


**Figure S5:** Distribution of evolutionary mechanisms across MIC trajectory classes. Trajectory arrow plots for each class of resistance gain. Arrow plots of Log2Fold change in mode ciprofloxacin MIC (μg/mL) from baseline against change in average maximal growth rate from baseline (ODhr-1). Each arrow traces the phenotypic change for a single patient, and follows time across the ORBIT-3 trial. Arrows are coloured by evolutionary mechanism for that patient.

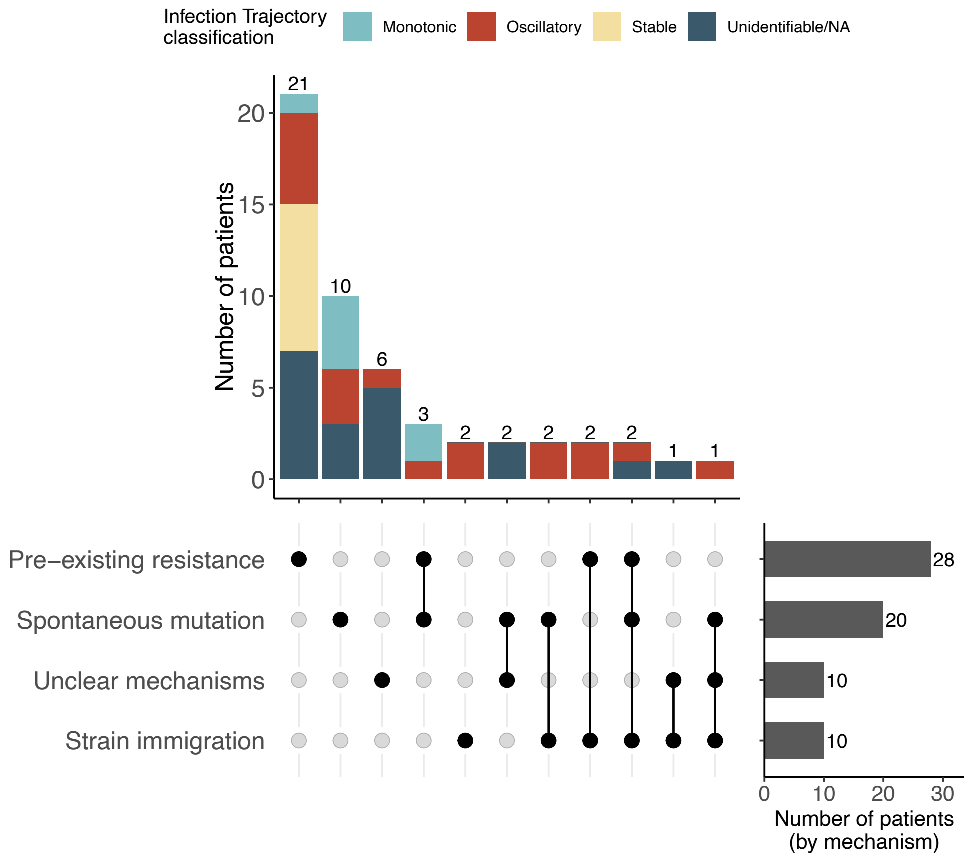


**Figure S6:** Distribution of evolutionary mechanisms present during ciprofloxacin treatment of infections for patients where resistance was detected at any timepoint. These mechanisms are present, but not necessarily responsible for driving emergence of resistance (i.e. they may occur after resistance was gained, or in the case of 2 patients, pre-existing resistance was present but then absent during all timecourse samples). UpSet matrix indicates each mechanism, with lines joining multi-mechanism combinations. Upper stacked bar plot indicates number of patients for each grouping, with colours indicating proportions of patients with each MIC trajectory classification (possibly for 33 patients). Right hand bar plot indicates total number of patients where each mechanism is present.

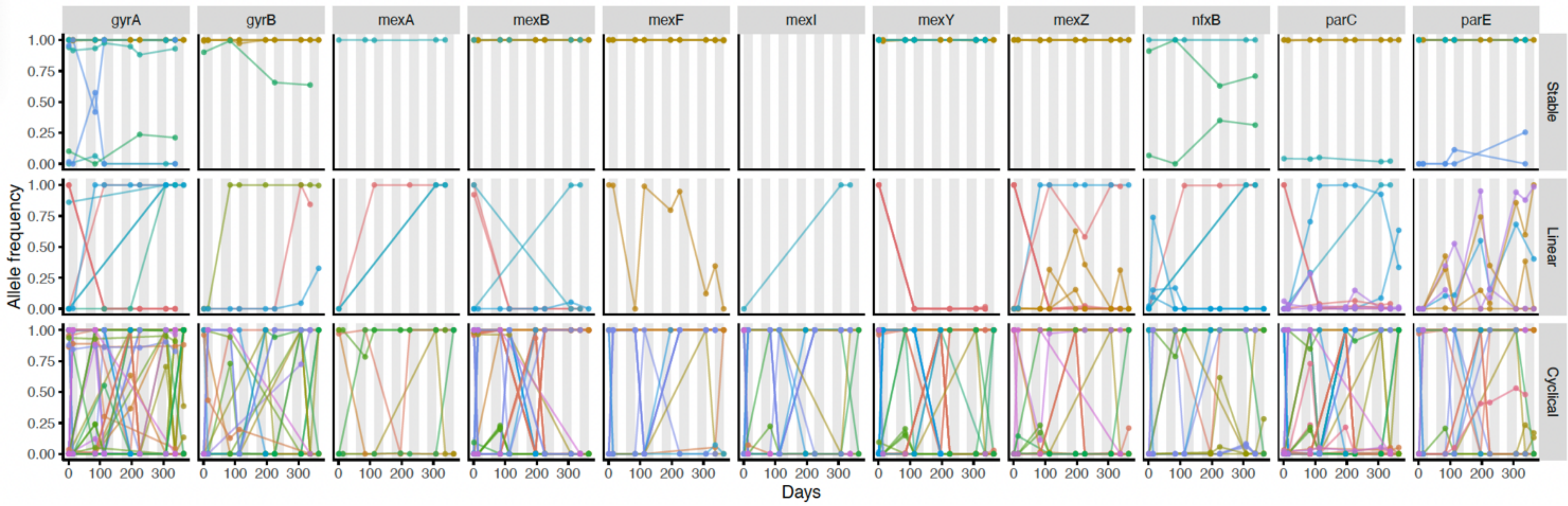


**Figure S7:** Infections followed distinct trajectories of ciprofloxacin resistance gain during the ORBIT-3 clinical trial. Allele frequency plotted against day of the trial. Lines represent individual mutations and are coloured by patient. Grey bars in plot area indicate ciprofloxacin dosing cycles (grey = on phase, white = off phase). Plots are faceted by gene and by phenotypic trajectory.

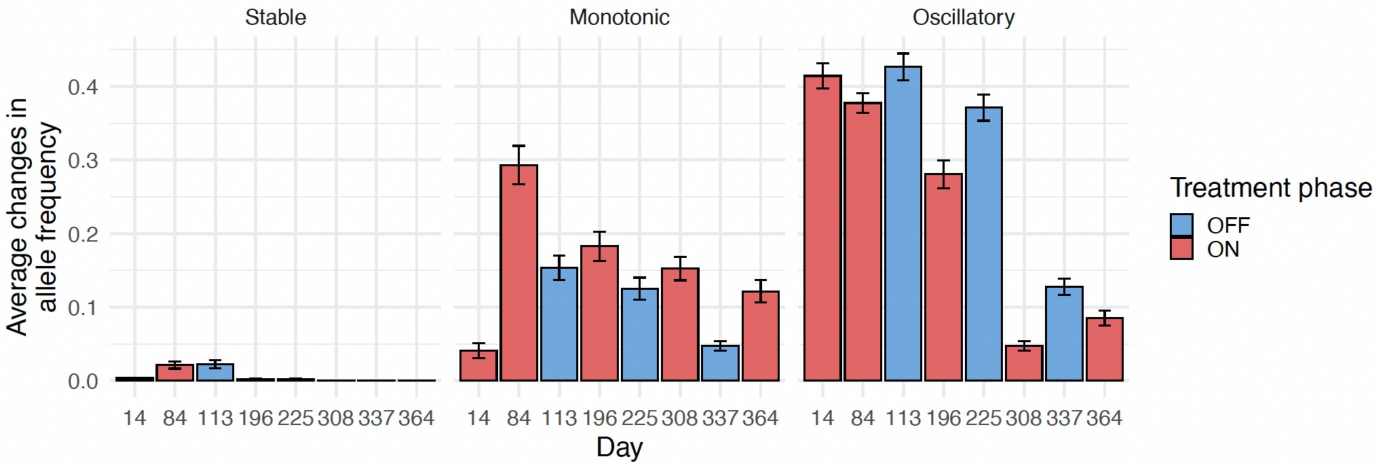


**Figure S8:** Average changes in allele frequency across the trial for each phenotypic trajectory class were calculated ignoring the direction of change. Each bar is calculated over the time period ending on the day indicated (except for Day 84, which was calculated from Day 1 on to keep it consistent with other on-off-on phases) and starting on the previous sampling day. Ciprofloxacin dosing indicated by colours: red shows on phases, blue off phases. Error bars indicate standard error.

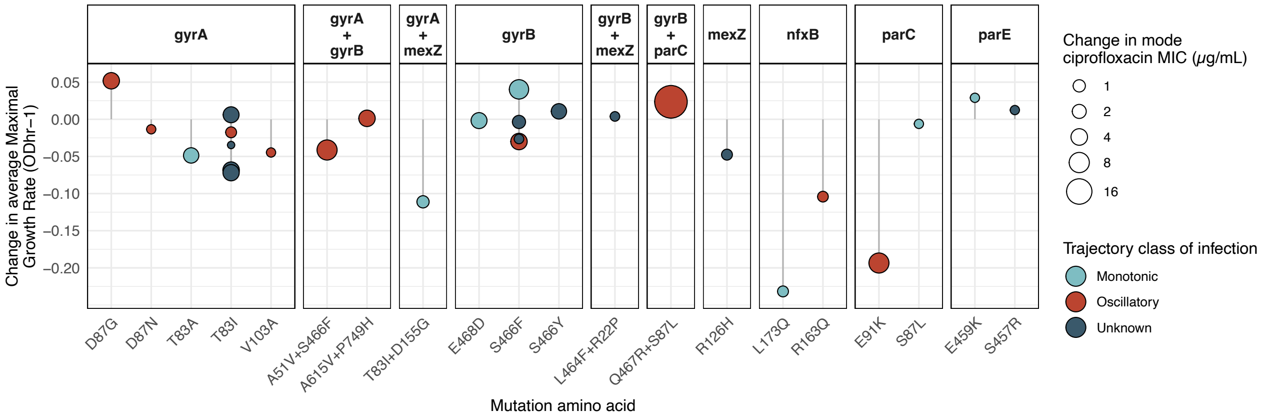
gyrA
gyrA + gyrB
gyrA + mexZ
gyrB
gyrB + mexZ
gyrB + parC
mexZ
nfxB
parC
parE
Change in average Maximal Growth Rate (ODhr−1)
0.05
0.00
−0.05
−0.10
−0.15
−0.20
D87G
D87N
T83A
T83I
V103A
A51V+S466F
A615V+P749H
T83I+D155G
E468D
S466F
S466Y
L464F+R22P
Q467R+S87L
R126H
L173Q
R163Q
E91K
S87L
E459K
S457R
Mutation amino acid
Change in mode ciprofloxacin MIC (μg/mL)
1
2
4
8
16
Trajectory class of infection
Monotonic
Oscillatory
Unknown

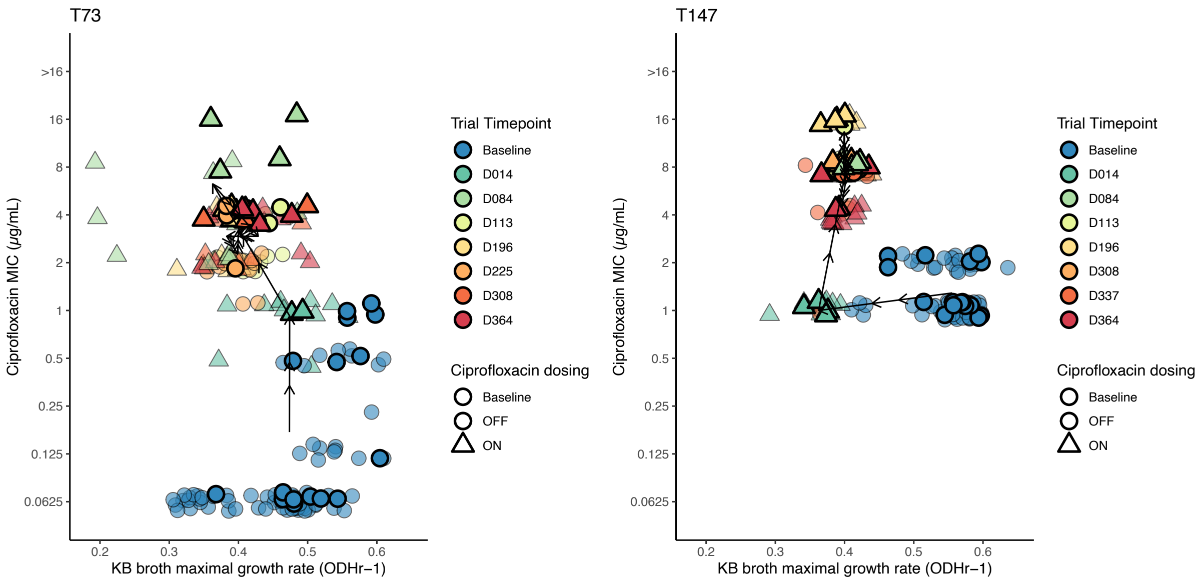
T73
T147
Ciprofloxacin MIC (μg/mL)
>16
16
8
4
2
1
0.5
0.25
0.125
0.0625
0.2
0.3
0.4
0.5
0.6
KB broth maximal growth rate (ODHr−1)
Trial Timepoint
Baseline
D014
D084
D113
D196
D225
D308
D337
D364
Ciprofloxacin dosing
Baseline
OFF
ON

A

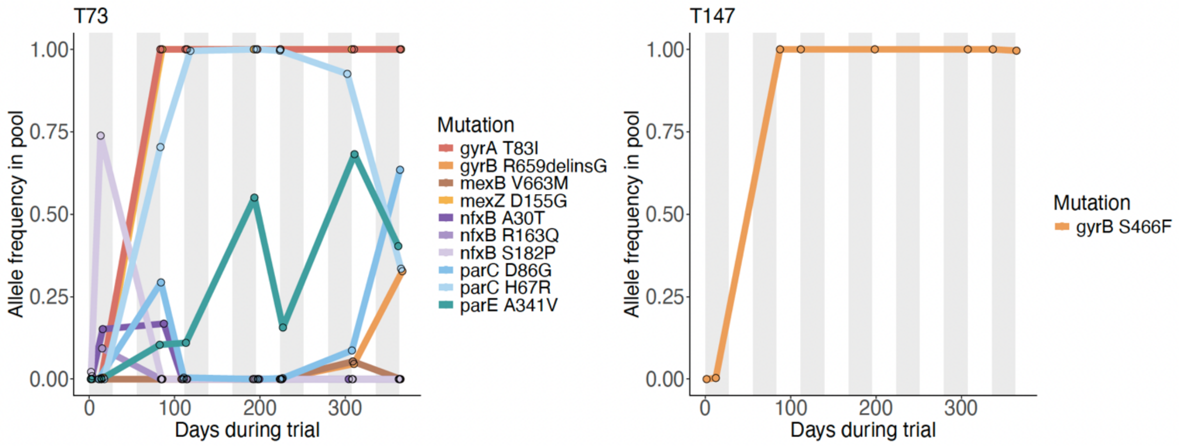
T73
T147
Allele frequency in pool
1.00
0.75
0.50
0.25
0.00
0
100
200
300
Days during trial
Mutation
gyrA T83I
gyrB R659delinsG
mexB V663M
mexZ D155G
nfxB A30T
nfxB R163Q
nfxB S182P
parC D86G
parC H67R
parE A341V
gyrB S466F

**Figure S9:** Emergence of ciprofloxacin resistance is frequently associated with costs to growth rate, which can persist for the trial duration. **A)** Change in average maximal growth rate (ODhr-1) upon gain of spontaneous mutations. Mutant genes are detailed above each facet, and mutation amino acid changes given on the x axis. Points coloured by trajectory classification of the infection they occur within, and scaled by change in the mode ciprofloxacin MIC (μg/mL) associated with the SNP. **B)** Phenotypic trajectory plot for patients T73 and T147. Data point colours indicate trial timepoints, shapes of points indicate ciprofloxacin dosing at each timepoint. The black arrow traces the mean values of each variable, with arrow direction indicating the passage of time across the trial. **C)** Allele frequency in pooled sample sequences for patients T73 and T147.
